# The insert sequence in SARS-CoV-2 enhances spike protein cleavage by TMPRSS

**DOI:** 10.1101/2020.02.08.926006

**Authors:** Tong Meng, Hao Cao, Hao Zhang, Zijian Kang, Da Xu, Haiyi Gong, Jing Wang, Zifu Li, Xingang Cui, Huji Xu, Haifeng Wei, Xiuwu Pan, Rongrong Zhu, Jianru Xiao, Wang Zhou, Liming Cheng, Jianmin Liu

## Abstract

At the end of 2019, the SARS-CoV-2 induces an ongoing outbreak of pneumonia in China^1^, even more spread than SARS-CoV infection^2^. The entry of SARS-CoV into host cells mainly depends on the cell receptor (ACE2) recognition and spike protein cleavage-induced cell membrane fusion^3,4^. The spike protein of SARS-CoV-2 also binds to ACE2 with a similar affinity, whereas its spike protein cleavage remains unclear^5,6^. Here we show that an insertion sequence in the spike protein of SARS-CoV-2 enhances the cleavage efficiency, and besides pulmonary alveoli, intestinal and esophagus epithelium were also the target tissues of SARS-CoV-2. Compared with SARS-CoV, we found a SPRR insertion in the S1/S2 protease cleavage sites of SARS-CoV-2 spike protein increasing the cleavage efficiency by the protein sequence aligment and furin score calculation. Additionally, the insertion sequence facilitates the formation of an extended loop which was more suitable for protease recognition by the homology modeling and molicular docking. Furthermore, the single-cell transcriptomes identified that ACE2 and TMPRSSs are highly coexpressed in AT2 cells of lung, along with esophageal upper epithelial cells and absorptive enterocytes. Our results provide the bioinformatics evidence for the increased spike protein cleavage of SARS-CoV-2 and indicate its potential target cells.

## Introduction

At the end of 2019, a rising number of pneumonia patients with unknown pathogen emerged from Wuhan to nearly the entire China^7^. A novel coronavirus was isolated and based on its phylogeny, taxonomy and established practice, the Coronavirus Study Group (CSG) recognized it as a sister to severe acute respiratory syndrome coronaviruses (SARS-CoVs) and labeled it as severe acute respiratory syndrome coronavirus 2 (SARS-CoV-2)^1,8^. Although SARS-CoV-2 is generally less pathogenic than SARS-CoV and Middle East respiratory syndrome coronavirus (MERS-CoV), it has a relatively high transmissibility^9^.

With regard to human coronavirus, the transmissibility and infectivity is largely controlled by the spike (S) surface envelope protein^10^. Its surface unit (S1) mediates the entry into host cells by binding to cell receptor and the transmembrane unit (S2) subunit regulates the fusion of viral and cellular membranes^3^. Prior to membrane fusion, the S protein should be cleaved and activated to allow for the fusion peptide releasing onto host cell membranes (Fig. 1a)^11^. SARS-CoV-2 uses the same cell receptor (angiotensin converting enzyme II, ACE2) as SARS-CoV, with a similar binding affinity, whereas their transmissibility and infectivity are different^5,6,12,13^ Thus, the different virus transmission and infectivity may be associated with the differentiated protease-induced S protein cleavage between SARS-CoV-2 and SARS-CoV.

**Fig. 1.**
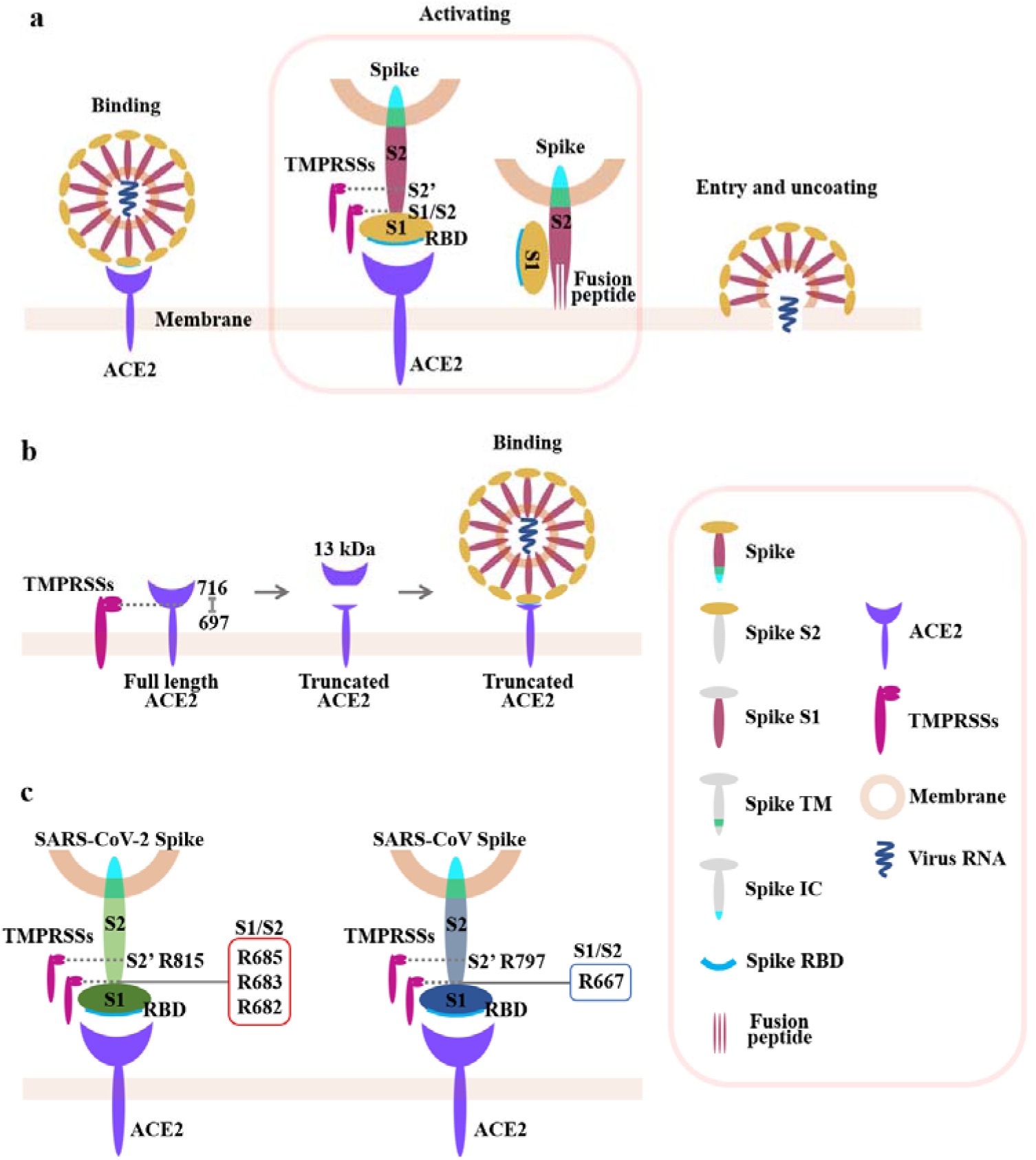
The schematic diagram of the project. a. The entry of SARS-CoV into host cells: The spike protein of SARS-CoV binds to ACE2 through its S1 subunit for viral recognition. Then it is cleaved by TMPRSS2 at the S1/S2 boundary or within S2 subunit, which removes the structural constraint of S1 on S2, and releases the internal fusion peptide combined with the Spike TM domain for the fusion of viral and cellular membranes. Finally, the viral genomes enter into the host cells. b. ACE2 cleaving by TMPRSSs: TMPRSS2 can also cleave ACE2 amino acids 697 to 716, resulting in the shedding of 13kD ACE2 fragment in culture supernatants and augmented viral infectivity. c. The difference between SARS-CoV-2 and SARS-CoV in the Spike protein cleavage: The Spike protein of SARS involves two cleavage sites recognized by TMPRSSs, one at arginine 667 and the other at arginine 797 (right). Compared with SARS-CoV, the Spike protein of SARS-CoV-2 (left) has an insertion sequence 680-SPRR-683 at the S1/S2 cleavage site. We speculated that R682, R683 and R685 (red box) could be used as the most suitable substrates for TMPRSSs, which can increase the Spike protein cleavage efficiency of TMPRSSs, promote its activation and enhance SARS-CoV-2 infection.

The transmembrane serine proteases (TMPRSSs) were the main host cell proteases on the cell membrane^14^. The substrate specificity of TMPRSSs are almost similar and revealing a strong preference for arginine or lysine residues in the P1 position. Nowadays, their hydrolytic effects of TMPRSSs have been widely reported in SARS-CoV and MERS-CoV pneumonia^15^. In the SARS-CoV-infected alveolar cells, TMPRSSs, especially the TMPRSS2 and TMPRSS11D, cleave the SARS-CoV S protein (SARS-S) at residue R667 (the S1/S2 cleavage site) and residue R797 (the S2’ cleavage site) (Fig. 1a)^15,16^. Besides cleaving S protein, they can also promote viral spread in the host by cleaving ACE2 (Fig. 1b)^14,17^. Although SARS-CoV-2 and SARS-CoV share the same host cell receptor with a similar affinity, however, the SARS-CoV-2 S protein cleavage induced by TMPRSS remains unclear which may be associated with the viral infectivity^4,5^.

## Results

### The comparison of the S1/S2 and S2’ cleavage sites between SARS-CoV-2 and SARS-CoV

Generally, compared with SARS-CoV, the major differences in SARS-CoV-2 are the three short insertions in the N-terminal domain and four out of five key residues changes in the receptor-binding motif^5^. Here we used the alignment, furin score and homology modeling to compare the sequence of the S1/S2 and S2’ cleavage sites between SARS-CoV-2 and SARS-CoV (Fig. 1c). The amino acid sequence of the S1/S2 and S2’ cleavage sites among ten beta-coronavirus were then analyzed and we found that compared with SARS, there was an insertion sequence (SPRR) in the S1/S2 cleavage sites of SARS-CoV-2 (Fig. 2a). The furin score was next used to identify the cleavage efficiency of the insertion sequence in SARS-CoV-2. Its furin score was 0.688, which was obviously higher than that of the corresponding sequence in SARS-CoV (0.139), indicating that the insertion sequence may increase the cleavage efficiency by proteases (Fig. 2b).

**Fig. 2.**
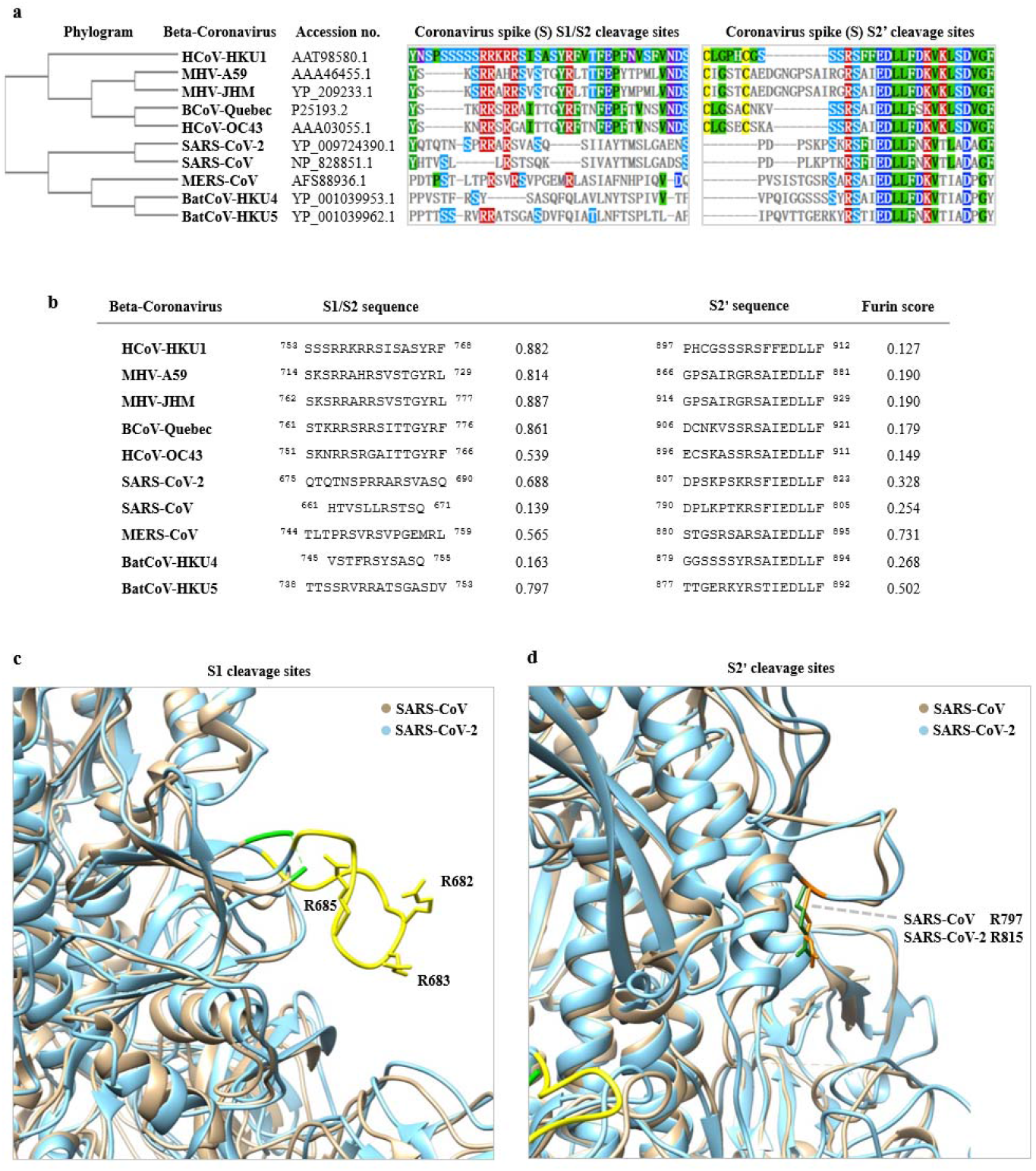
The two potential Spike protein cleavage sites of SARS-CoV and SARS-CoV-2 by TMPRSS2. a. Phylogenetic tree based on the protein sequences of Spike protein in SARS-CoV-2, SARS-CoV and other eight beta-coronaviruses are presented, along with the amino acid sequence alignment of two potential cleavage sites by TMPRSS2. b. The putative furin scores of the two potential cleavage sites of the ten coronaviruses. c. Structure comparison of the detailed Spike protein of the SARS-CoV and SARS-CoV-2. The insert 675-690 of SARS-CoV-2 Spike protein (yellow) and the corresponding loci to SARS-CoV Spike protein 661-672 (green). Three important residues, R682, R683, R685, are specially marked. d. The detail of c. The similarly SARS-CoV R797 with SARS-CoV-2 R815 are marked with forest green and orange, respectively.

The structures of SARS-S and SARS-CoV-2 S protein were presented in Extended Data Fig. 1a and 1b, along with their structural superimposition (Extended Data Fig. 1c). The structural comparison of homology modeling SARS-CoV-2 S protein with SARS-S protein (PDB: 5×5b) showed that a exposed loop was formed by the insertion which comprised R682 and R683 (S1/S2 site) on the surface of SARS-CoV-2 S protein, and no significant difference of them in S2’ site (Fig. 2c, d).

### The insertion sequence of SARS-CoV-2 facilitating the TMPRSS recognition and S protein cleavage

Structurally, TMPRSSs include extracellular domain, transmembrane domain and intracellular domain in which extracellular domain is the main catalytic domain. They show similar substrate-specificity and catalytic mechanism. Take TMPRSS2 as an example. The catalytic triad consisted of H296, D345 and S441 and the substrate binding residue D435, a conserved aspartate residue, was located in the bottom of pocket^18,19^. The substrate binding pocket is deeper than most of serine proteinase (Extended Data Fig. 2a, b). The bottom of the catalytic pocket has a negatively charged aspartic acid residue which can facilitate the binding and stabilization of arginine or lysine residues in the P1 position^18,19^.

Polypeptide substrate analogue KQLR included arginine, glutamine, leucine and lysine (Extended Data Fig. 2c). The substrate analogue could bind to the catalytic pocket of TMPRSS2 (Extended Data Fig. 2d, e). The conformation of the insertion sequence in SARS-CoV-2 S protein and TMPRSS2 was next simulated by molecular docking. We found the insertion sequence formed a loop which was easily recognized by the catalytic pocket of TMPRSS2 (Extended Data Fig. 2f, g). Thus, both the furin score and molecular docking revealed that the insertion sequence of SARS-CoV-2 facilitates the TMPRSS2 recognition and S protein cleavage.

### The potential target tissues of COVID-19

The entry of SARS-CoV-2 into host cells depends on the cell receptor recognition and cell proteases cleaving. Thus, the target cells should coexpress both the cell receptor ACE2 and cell proteases TMPRSSs. In order to identify the coexpressing cell composition and proportion, we utilized 3 datasets including 32 samples and built the largest single-cell transcriptome atlas of normal lung, the commonest infected organ of SARS-CoV-2.

After initial quality controls, a total of 113,045 cells and 29 sub-clusters were identified in the lung (Fig. 3a). The marker genes and dataset proportions of each sub-cluster were presented in Extended Data Fig. 3–4.

**Fig. 3.**
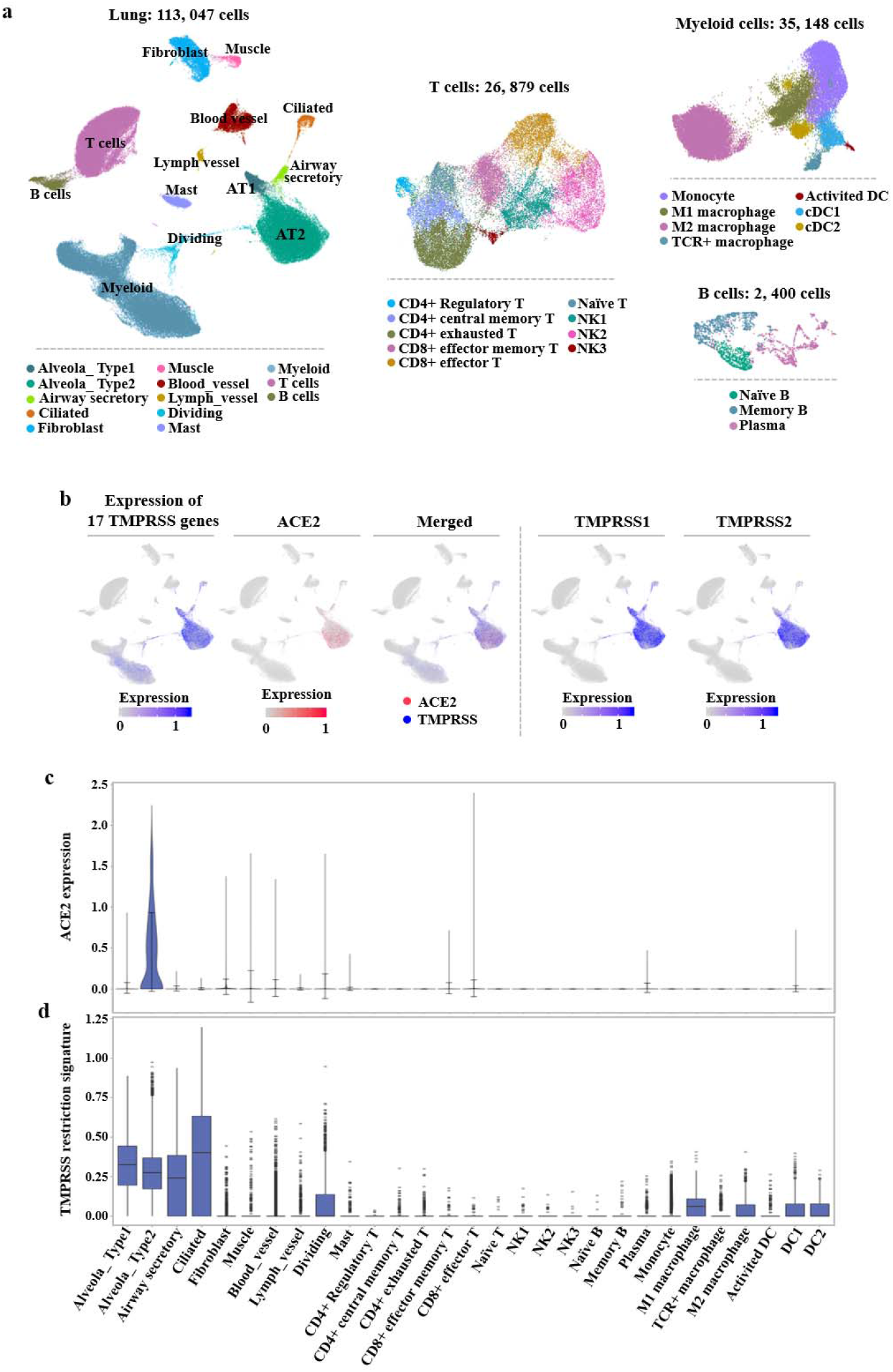
Single-cell analysis of the normal lung tissue. a. The UMAP plots of the landscape of lung cells. Thirteen clusters are colored, distinctively labeled. T, B and myeloid cell subsets are further divided into finer cell subsets according to the heterogeneity within the cell population. b. The feature plots of the 17 TMPRSS genes, ACE2, TMPRSS1 and TMPRSS2. c. The expression of ACE2 across clusters in the violin plot. The expression is measured as the log2 (TP10K+1) value. d. The mean expression of TMPRSS family genes across clusters in the boxplot. The expression is measured as the mean log2 (TP10K+1) value.

We detected the expression of ACE2 and TMPRSSs in 29 cell groups, in which the expression of the whole 17 TMPRSS genes is in the form of total signature value. Pseudodyeing analysis was performed and we found that ACE2 was mainly expressed in AT2 cells and marked with red (Fig. 3b, c). The total 17 TMPRSS genes was found in AT1, AT2, airway secretory and ciliated cells colored with blue (Fig. 3b, d, Extended Data Fig. 5a). Thus, we found an obvious coexpression between TMPRSSs and ACE2 in AT2. Among the whole TMPRSS genes, TMPRSS1 and TMPRSS2 were highly expressed in AT2 and AT1 cells, which were co-expressed with ACE2 in lung (Fig. 3b, Extended Data Fig. 5b). Due to the entry of virus into host cell is related to endocytosis, we also detected the endocytosis-related genes among different cells. We found that these genes had consistent distribution and highly expressed in AT1, AT2, airway secretory, ciliated cells and M2 macrophage (Extended Data Fig. 5c).

Due to the RNA of SARS-CoV-2 was also found in the stool specimen of the SARS-CoV-2-infected patient^20^, the digestive system may also be the potential route of COVID-19. Thus, in addition to lung, 4 datasets with the single-cell transcriptomes of the esophagus, gastric, small intestine and colon were analyzed to identify the expression of ACE2 and TMPRSSs in the digestive system. The co-expression of

ACE2 and TMPRSS was analyzed in esophagus, stomach, small intestine and colon by 87947, 29678, 11218 and 47442 high-quality single cells, respectively (Extended Data Fig. 6a). The coexpression of ACE2 and total TMPRSS genes were found in the upper epithelial cells of esophagus, the absorptive enterocytes of ileum epithelia and the enterocytes of colon epithelia (Extended Data Fig. 6b-e, 7a-d).

As both ACE2 and TMPRSSs are expressed in the lung and digestive system, we next compared their relative expression values in the ACE2-expressing cells. A similar distribution was found between ACE2 and TMPRSSs in all the 9 clusters with high expressions in the esophageal upper epithelial cells, the ileal absorptive enterocytes and the colonic enterocytes (Fig. 4a). In addition, their expression of AT2 was relatively lower than that of epithelial cells in the digestive system. Among all the TMPRSSs, TMPRSS1 and TMPRSS2 were relatively highly expressed in AT2, and most TMPRSSs were highly found in the esophageal upper epithelial cells (Extended Data Fig. 8a). The endocytosis- and exocytosis-associated genes which are related to the entry of virus into host cells and virus infection were also detected in all the 9 clusters. The endocytosis signature was more expressed in AT1 and AT2 cells, whereas the exocytosis signature was highly gathered in esophageal upper epithelial cells. It can explain that the commonest infected tissue in COVID-19 is pulmonary alveoli and SARS-CoV-2 can also be detected in the esophageal erosion (Fig. 4b)^21^. The RNA-seq data of lung, esophagus, stomach, small intestine, colon-transverse and colon-sigmoid were obtained from GTEx database. The expressions of ACE2 and TMPRSS2 also had a similar tendency and were highly expressed in small intestine and colon, while the TMPRSS11D was mainly found in the esophagus (Extended Data Fig. 8b).

**Fig. 4.**
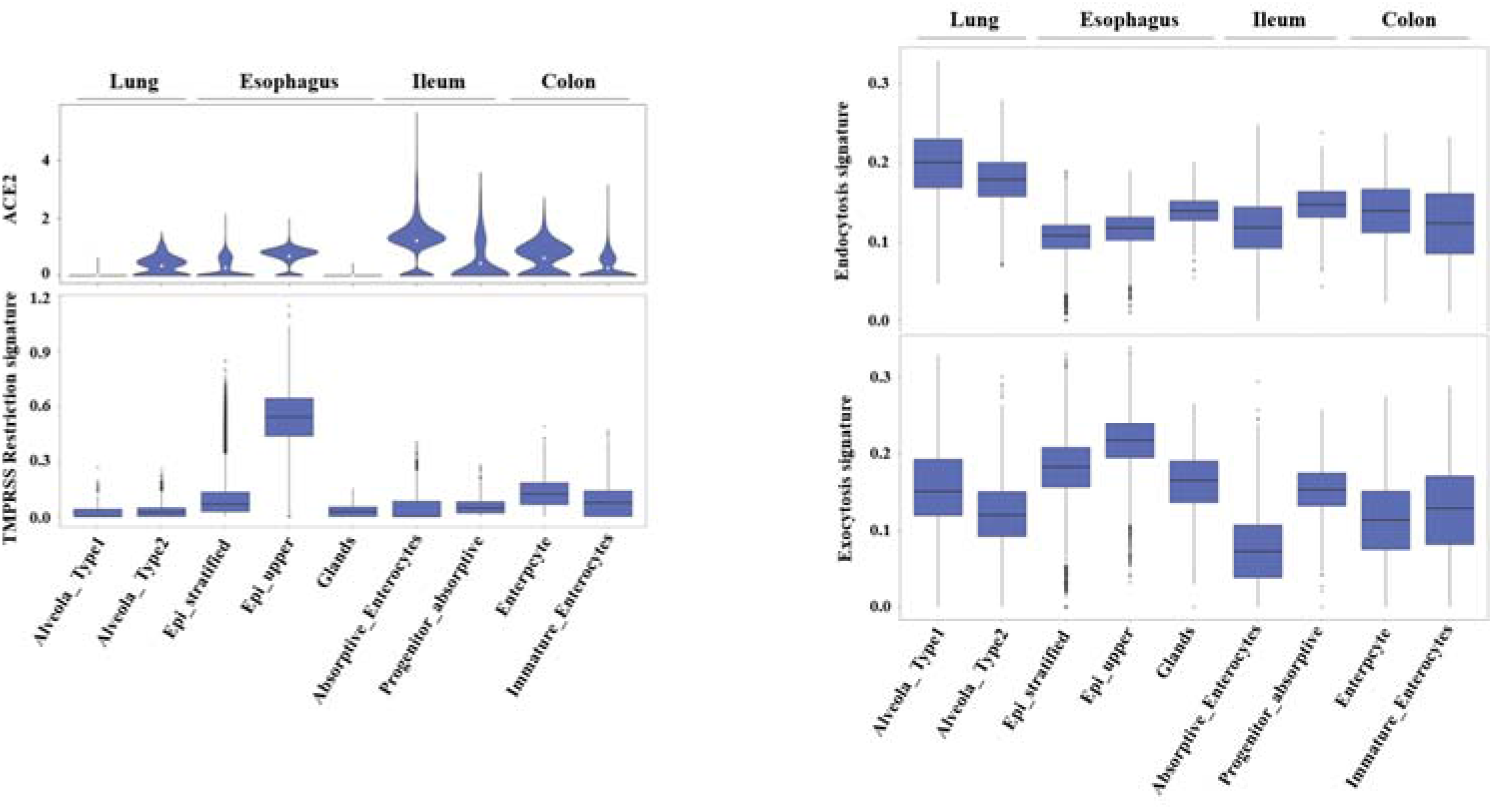
Expression levels of ACE2, TMPRSS restriction signature and functional gene sets in lung and digestive tracts. a. The expression levels of ACE2 and TMPRSS restriction signature in 2 lung clusters and 7 digestive tract clusters. The expression is measured as the log2 (TP10K+1) value. b. The expression levels of endocytosis and exocytosis-associated genes in 2 lung clusters and 7 digestive tract clusters. The expression is measured as the log2 (TP10K+1) value.

## Discussion

The coronaviruses is the common infection source of respiratory, enteric and central nervous system in humans and other mammals^22^. At the beginning of the twenty-first century, two betacoronaviruses, SARS-CoV and MERS-CoV, result in persistent public panics and became the most significant public health events^23^. In December 2019, a novel identified coronavirus (SARS-CoV-2) induced an ongoing outbreak of pneumonia in Wuhan, Hubei, China^7^. The rapidly increasing number of SARS-CoV-2-infected cases suggests that SARS-CoV-2 may be transmitted effectively among humans and give rise to a high pandemic potential^7,8,24^ Previous studies identified that SARS-CoV mutated between 2002 and 2004 to better bind to its cell receptor, replicate in human cells and enhance the virulence^9^. Thus, it is important to explore whether SARS-CoV-2 behaves like SARS-CoV to adapt to the host cell. Notably, SARS-CoV and SARS-CoV-2 share the same receptor protein ACE2^5,25^. Besides, the receptor-binding domain (RBD) in S protein of SARS-CoV-2 binds to ACE2 with the similar affinity as SARS-CoV RBD does^6^. Thus, during the process of viral and host cellular membrane fusion, whether the specific structure of SARS-CoV-2 S protein seems better suited to be activated by host cell proteases may be related to the different virus infectivities and transmissibilities between SARS-CoV and SARS-CoV-2^6^.

In this study, we found the furin score of the S1/S2 cleavage sites in SARS-CoV-2 was higher than that of SARS, implying a more degree of hydrolysis. Through the comparison of the two structures, R682, R683 and relative S680, P681 extended the original exposed loop combined with R685 of SARS-CoV-2, which was more suitable for hydrolysis by TMPRSSs. The substrate specificity of TMPRSSs are almost similar, revealing a strong preference for arginine or lysine residues in the P1 position represented by R. More R (R682, R683 and R685) in the S1/S2 cleavage sites of SARS-CoV-2 can enhance the cleavage of S1 with S2, which means that the structurally constrains of S1 on S2 is removed, and the fusion peptides in S2 are exposed and insert into the target host cell membrane, finally it increases the efficiency of fuse membranes^18,19^.

By the way, some researchers previously supposed the SARS-CoV-2 was artificial due to four inserts in the S protein of SARS-CoV-2 from HIV sequence. However, the results of protein sequence alignment revealed that the similar sequence of the reported fourth insertion site (680-SPRR-683) in SARS-CoV-2 was commonly found in many beta-coronavirus. Therefore, we supposed that based on the current evidence, it is not scientific to consider the insertion sequence in SARS-CoV-2 S protein being artificial.

With the help of single cell sequencing, we found a strong co-expression between ACE2 and TMPRSSs, in especial TMPRSS1and TMPRSS2, in lung AT2 cells, which was also the main infected cell type in SARS-CoV pneumonia^26^. Moreover, we also found the endocytosis-associated genes was highly expressed in AT2 cells, implying that endocytosis may also facilitate the entry of SARS-CoV-2 into host cells. As the alveolar stem-like cells, AT2 cells are in charge of surfactant biosynthesis, self-renewal and immunoregulation^27^. Thus, SARS-CoV-2 not only damages the AT2 cells leading to the direct injury to alveoli, but also raises alveolar surface tension to induce dyspnea^28^. Additionally, the injuryed AT2 also damages the immunologic balance in alveoli and results in inflammatory cascade^29^. In addition, they are also highly co-expressed in absorptive enterocytes and upper epithelial cells of esophagus, implying that intestinal epithelium and esophagus epithelium may also be the potential target tissues. This can explain the cases whose SARS-CoV-2 was detected in the esophageal erosion or stool specimen, implying that the digestive system is a potential route of COVID-19^7,20,21^.

Due to the critical role of TMPRSSs in influenza virus and coronavirus infections, serine protease inhibitors, such as camostat, nafamostat and leupeptin, have been used in the antiviral treatment targeting TMPRSSs with high antiviral activities^14,30,31^. Nowadays, Remdesivir (GS-5734) has been used in the treatment of SARS-CoV-2, however, the therapeutic effects are still unclear. Based on our results, we also supposed that TMPRSSs may also serve as candidate antiviral targets for SARS-CoV-2 infection and the clinical trials of serine protease inhibitors should also be performed for COVID-19.

## Methods

### Structure modelling

The structures of SARS-CoV-2 S protein and TMPRSS2 were generated by SWISS-MODEL online server^32^. The structures were marked, superimposed and visualized by Chimera^33^. To further explore the possible catalytic mechanism of the SARS-CoV-2 S protein cleaved by TMPRSS2, ZDOCK program was used to predict their interaction^34^ A total of 5000 models were generated and were set to 50 clusters, then the best scoring models from the 5 largest clusters were selected for further analysis.

### Furin score

The fragmentation maps, scoring and residue coverage analysis were conducted using arginine and lysine propeptide cleavage sites prediction algorithms ProP 1.0 server^35^.

### Single cell transcriptome data sources

Single cell transcriptome data were obtained from Single Cell Portal (https://singlecell.broadinstitute.org/single_cell), Human Cell Atlas Data Protal (https://data.humancellatlas.org) and Gene Expression Omnibus (GEO; https://www.ncbi.nlm.nih.gov/). Esophageal and lung data were obtained from the research of E Madissoon *et al* containing 21 esophageal and 19 lung tissue samples^36^. Two lung datasets were further obtained from GSE122960^38^ and GSE128169^39^, including eight and five lung tissues respectively. GSE134520 included 6 gastric mucosal samples from 3 non-atrophic gastritis and 2 chronic atrophic gastritis patients^40^. GSE134809 comprises 11 noninflammatory ileal samples from Crohn’s disease patients^41^. The data from Christopher S *et al* consisted of 12 normal colon samples^42^.

### Quality control

Cells would be identified as poor-quality once (1) the number of expressed genes fewer than 200 or greater than 5000, or (2) more than 20% of UMIs being mapped to mitochondrial or ribosomal genes.

### Data Integration, Dimension Reduction and Cell Clustering

Different methods were performed to process the downloaded data:

1. Esophagus dataset. Rdata were obtained and dimension reduction and clustering had already been implemented by the authors^36^.
2. Lung, stomach and ileum datasets. We utilized functions in the Seurat package to normalize and scale the single-cell gene expression data^43^. Unique molecularidentifier (UMI) counts were normalized by the total number of UMIs per cell, multiplied by 10000 for normalization and log-transformed using the NormalizeData’’ function. Then, multiple sample data within each dataset were merged using the “FindIntegrationAnchors” and “Integratedata” functions. After identifying highly variable genes (HVGs) using the “FindVariableGenes” function a principal component analysis (PCA) was performed on the single-cell expression matrix using the ‘‘RunPCA’’ function. The ‘‘FindClusters’’ function in the Seurat package was next utilized to conduct the cell clustering analysis into a graph structure in PCA space after constructing a K-nearest-neighbor graph based on the Euclidean distance in PCA space. Uniform Manifold Approximation and Projection (UMAP) visualization was performed for obtaining the clusters of cells.
3. Colon Dataset. The single cell data was processed with the R packages LIGER^44^ and Seurat^43^. The gene expression matrix was first normalized to remove differences in sequencing depth and capture efficiency among cells. Variable genes in each dataset were identified using the “selectGenes” function. Then we used the “optimizeALS” function in LIGER to perform the integrative nonnegative matrix factorization and selecte a k of 15 and lambda of 5.0 to obtain a plot of expected alignment. The “quantileAlignSNF” function was then performed to builds a shared factor neighborhood graph to jointly cluster cells, then quantile normalizes corresponding clusters. Next nonlinear dimensionality reduction was calculated using the “RunUMAP” function and the results were visualized with UMAP.

### Identification of cell types and Gene expression analysis

Clusters were annotated on the expression of known cell markers and the clustering information provided in the articles. Then, we utilized the “RunALRA” function to impute lost values in the gene expression matrix. The imputed gene expression was shown in Feature plots and violin plots. We used “Quantile normalization” in the R package preprocessCore (R package version 1.46.0. https://github.com/bmbolstad/preprocessCore) to remove unwanted technical variability across different datasets. The data were further denoised to compare the gene expression levels of gene signature.

Endocytosis or exocytosis associated genes were obtained from Harmonizome dataset^45^.Mean expressions of the genesets were calculated to compare the ability of endocytosis or exocytosis among clusters.

To minimize bias, external databases of Genotype-Tissue Expression (GTEx)^46^ was used to detect gene expression of ACE2, TMPRSS1 and TMPRSS2 at the tissue levels including normal lung and digestive system, such as esophagus, stomach, small intestine and colon.

## Acknowledgements

This study was jointly supported by the National Natural Science Foundation of China (Grants 81702659 and 81572746) and National Key R&D Program of China (Grants 2016YFA0100800).

## Author contributions

J.L., L.C., W.Z. and J.X. conceived the idea and directed the team. T.M., H.C., H.Z. and W.Z. designed and coordinated the analysis and characterization. H.Z., Z.K., D.X., H.G. performed single-cell sequencing and characterization under the guidance of X.C., H.X., and H.W.. Data collection and generation were performed by J.W., Z.L., R.Z. and X.P.. Data interpretation was performed by J.L., L.C., W.Z. and J.X.. The alignment and structure comparison was performed by H.C. under the guidance of W.Z. The manuscript was written by T.M., H.C., Z.K. and W.Z. All authors contributed to the analysis and discussion of the results leading to the manuscript.

---

**Extended Data Fig. 1.**
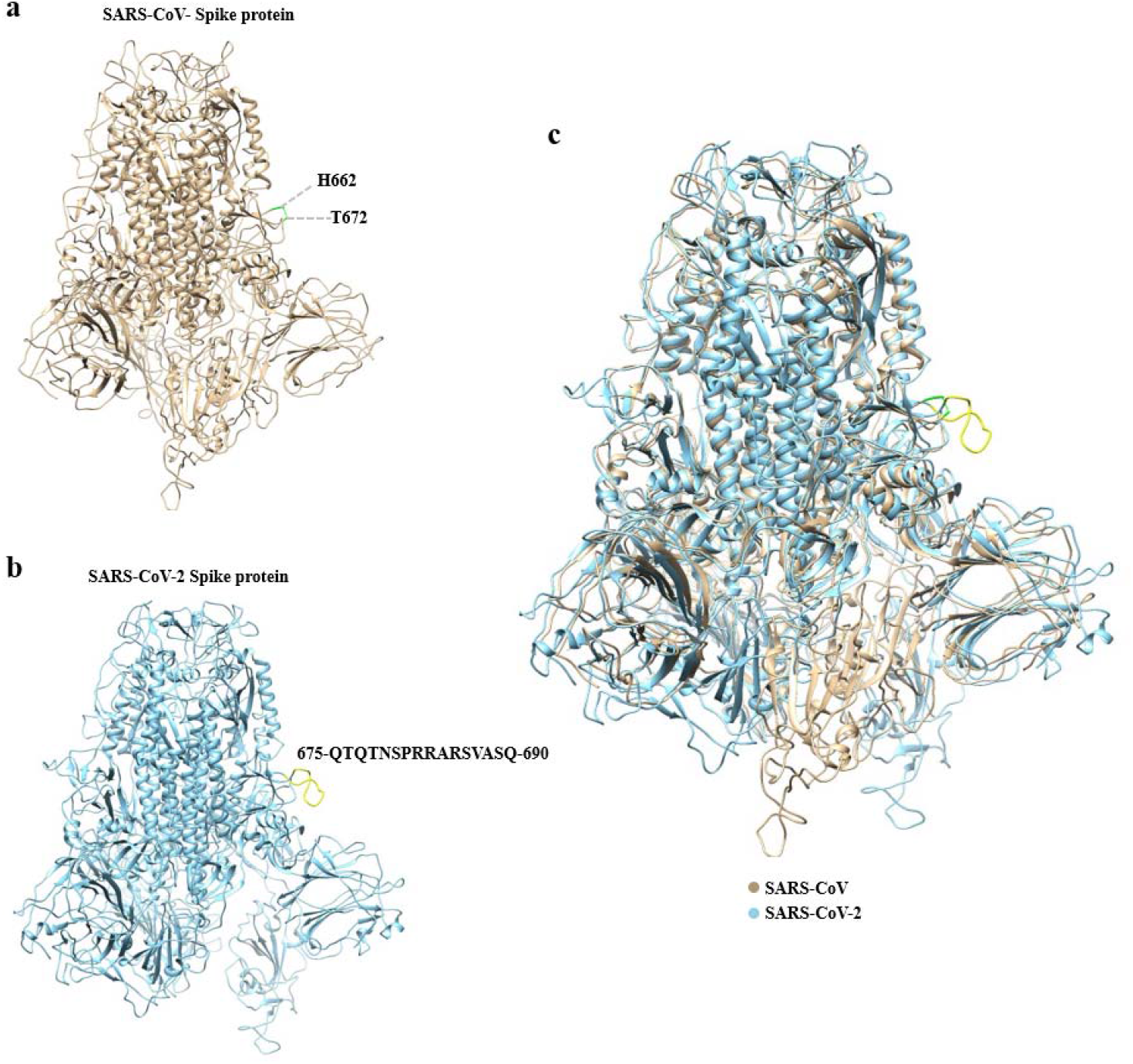
The overall structure of the Spike protein in SARS-CoV and SARS-CoV-2 homo-trimers. a. The structure of the SARS-CoV Spike protein (from PDB: 5X5B). The insert aa675-690 to SARS-CoV Spike protein aa661-672 with the structural missed residues are marked with green. b. The structure of the SARS-CoV-2 Spike protein (Modelled by SWISS-MODEL). The insert aa675-690 of 2019-nCoV Spike protein that corresponds to the insert region of SARS-V Spike protein is marked with yellow. c. The structural superimpose of Spike protein in the SARS-CoV (yellow) and SARS-CoV-2 (blue).

**Extended Data Fig. 2.**
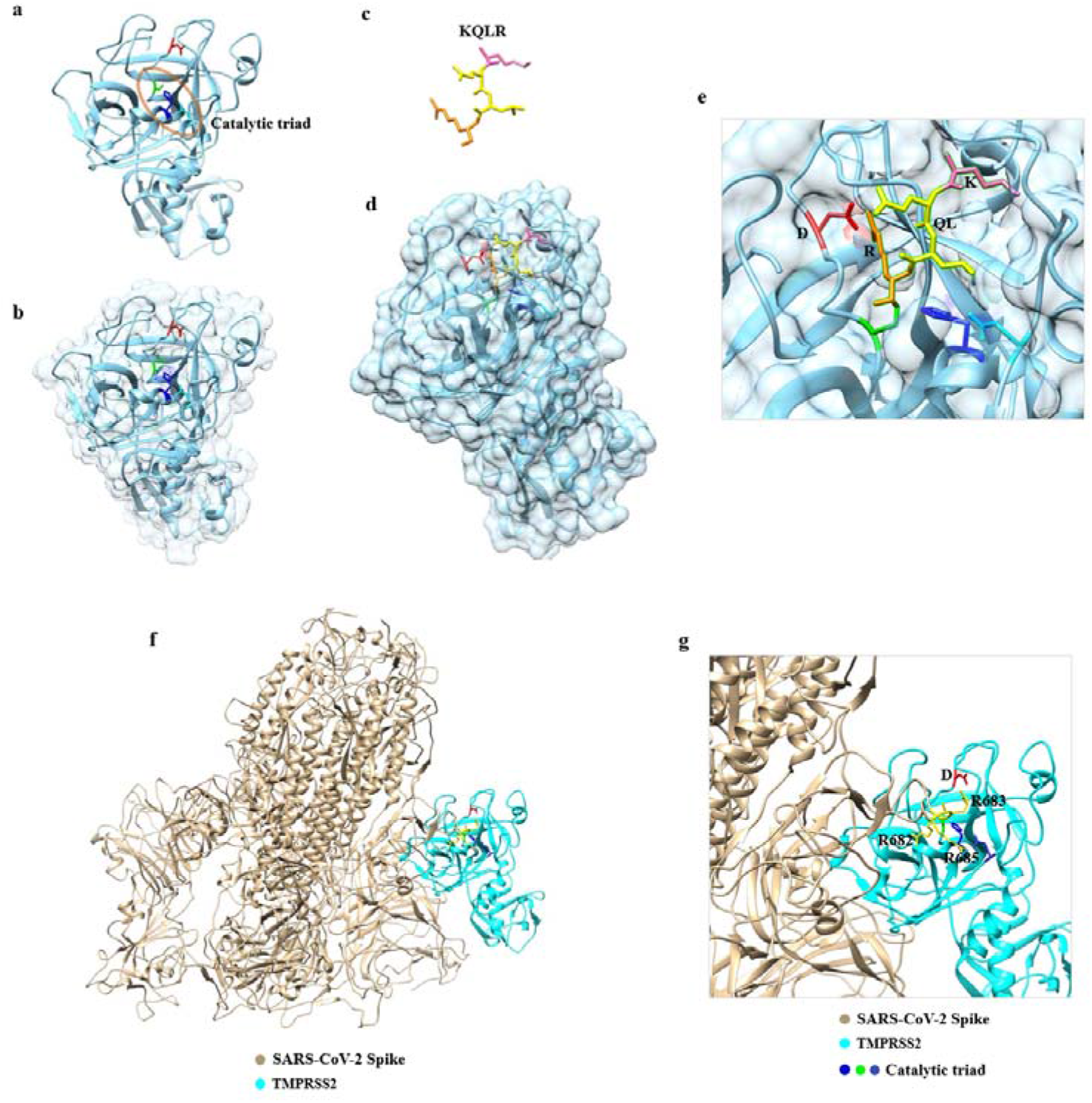
The structure and catalytic mechanism of TMPRSS2. a-b. The overall structure and surface of TMPRSS2 (Modelled by SWISS-MODEL). The TMPRSS2, catalytic triad comprised of H296, D345 and S441 are marked with cyan, blue, cyan and green, respectively. The substrate binding residue D435 located in the bottom of pocket is marked with red. c. The polypeptide substrate analogue KQLR. The cleavage site Arg is marked with orange. Gln and Leu are marked with yellow. Lys is marked with pink. d. The state of substrate analogue binding in the catalytic pocket. The state of substrate analogue binding in the catalytic pocket. e. The detail of d. Arg of substrate analogue is strongly interacted with D435 f. The predicted state of SARS-CoV-2 Spike protein binding to the catalytic pocket of TMPRSS2. g. The detail of f. SARS-CoV-2 Spike protein and D345 of TMPRSS2 are marked with wheat and medium blue, respectively.

**Extended Data Fig. 3.**
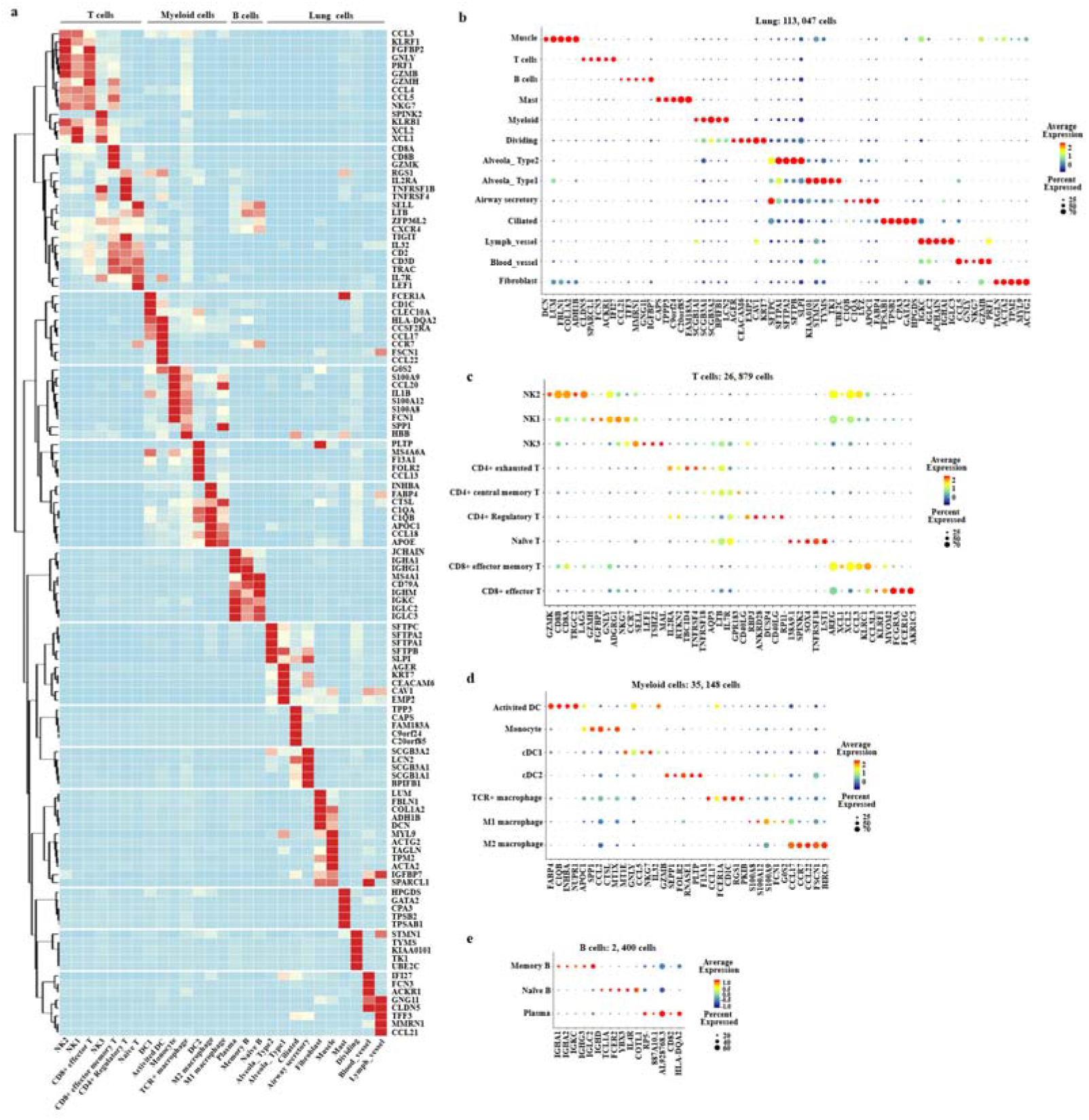
Subset-specific markers. a. The heatmap of marker genes (rows) across cell subsets (columns). The bubble diagram of marker genes in thirteen clusters (b) and the sub-clusters of T cells (c), B cells (d) and Myeloid cells (e).

**Extended Data Fig. 4.**
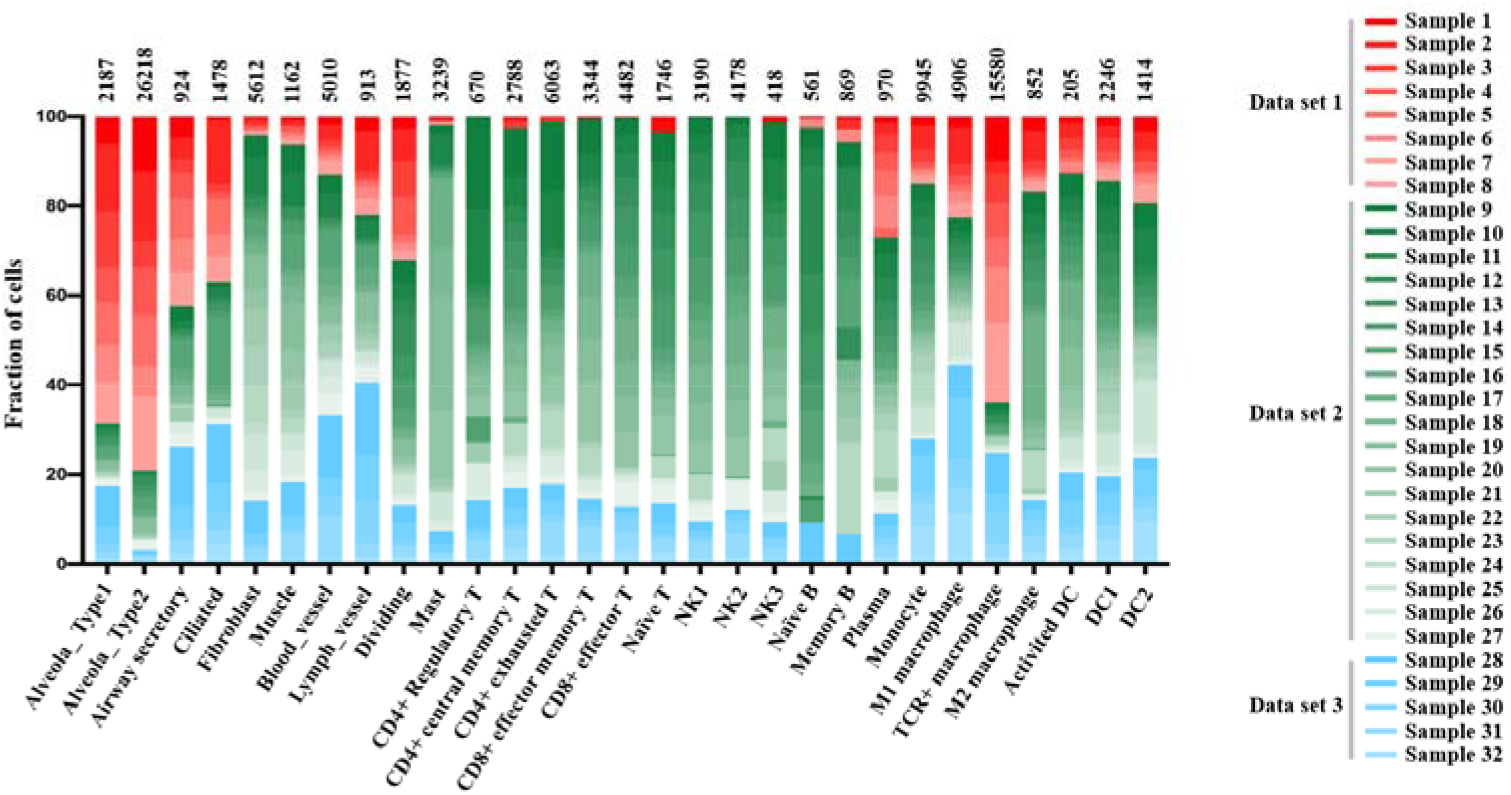
All cell subset distributions across samples. The fractions of cells (y axis) in each cell subset (bars) that are derived from each sample in 3 databases (red, green and blue). The numbers of cells in each cluster are labeled above.

**Extended Data Fig. 5.**
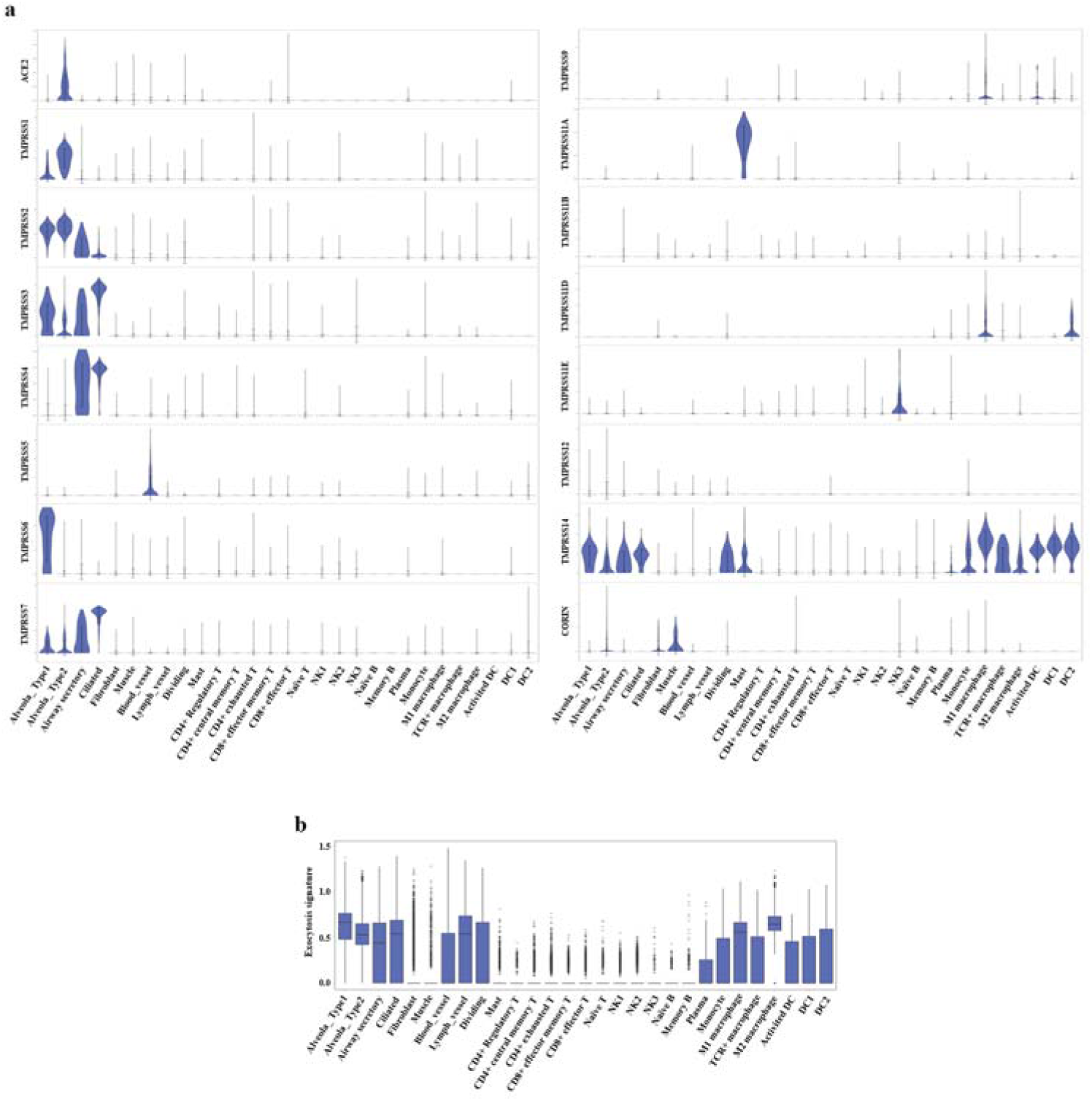
The expression levels of ACE2, TMPRSS genes and exocytosis-associated genes in lung subsets. a. The Violin plots of ACE2 and TMPRSS family genes across clusters. The expression is measured as the log2 (TP10K+1) value. b. The boxplot of exocytosis-associated across clusters. The expression is measured as the mean log2 (TP10K+1) value.

**Extended Data Fig. 6.**
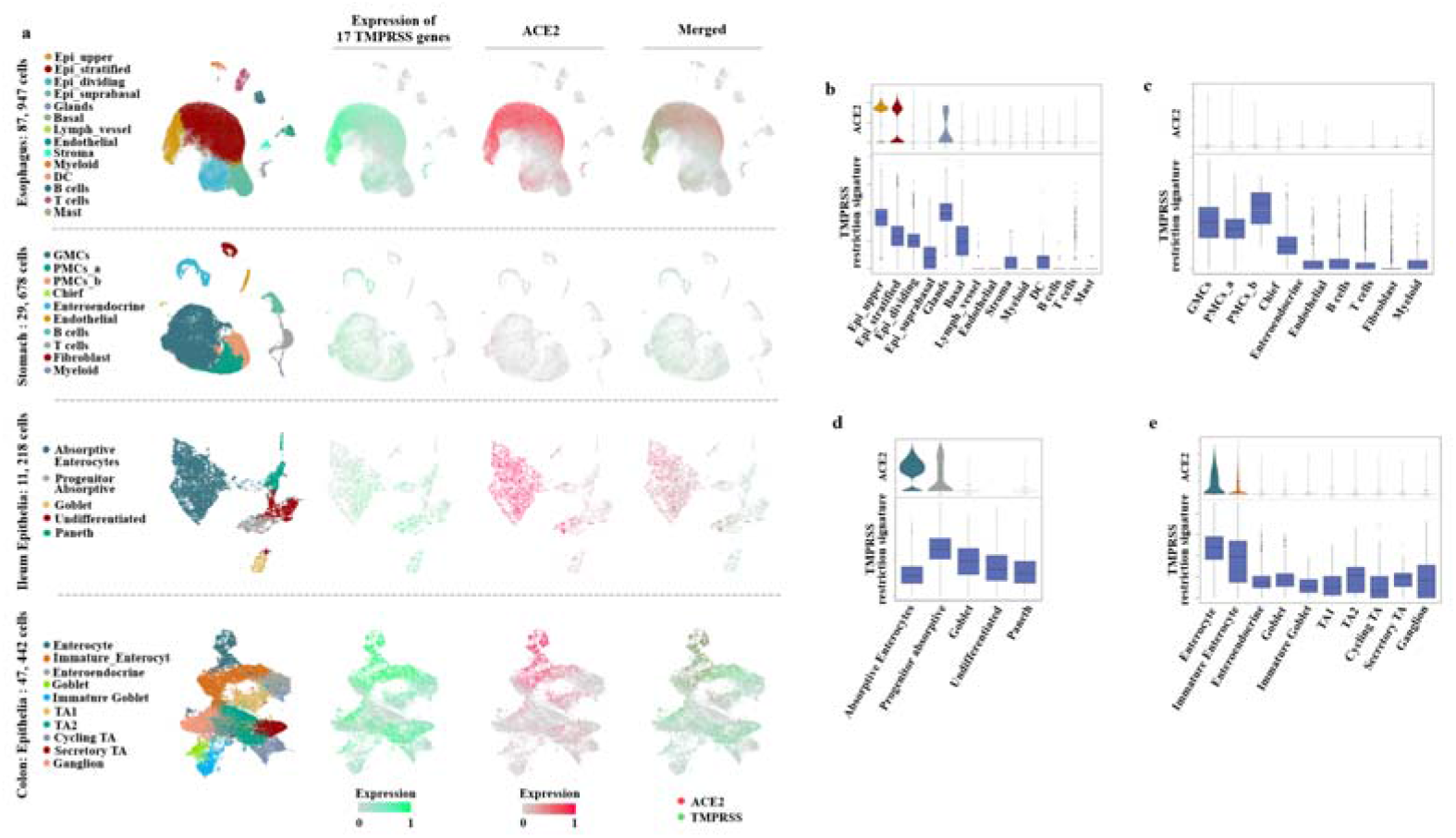
The single-cell analysis of esophageal cells, gastric mucosal cells, ileal epithelial cells and colonic epithelial cells. a. The UMAP plots of esophageal cells, gastric mucosal cells, ileal epithelial cells and colonic epithelial cells. The Feature plots show the expression of ACE2 (red) and TMPSS family genes (green). The plots were merged to reveal the co-expression of these genes (brown). c. The expression levels ACE2 and TMPRSS restriction signature across clusters in esophagus (b), stomach(c), ileum(d) and colon(d). The expression is measured as the mean log2 (TP10K+1) value.

**Extended Data Fig. 7.**
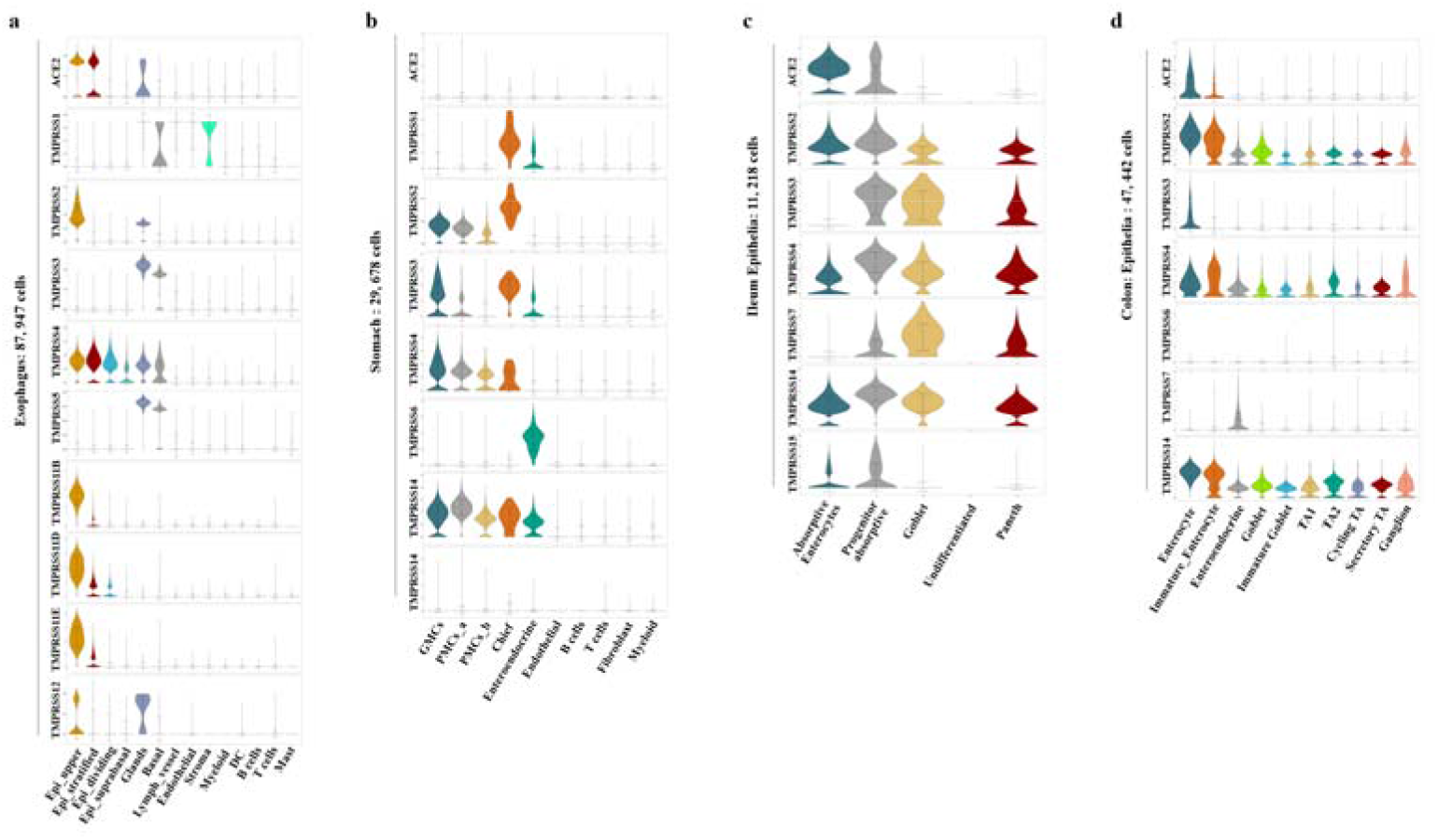
The expression levels of ACE2 and TMPRSS family genes in lung and digestive tracts. The violin plots of ACE2 and TMPRSS family genes across clusters in esophagus (a), stomach (b), ileum (c) and colon (d). The expression is measured as the mean log2 (TP10K+1) value.

**Extended Data Fig. 8.**
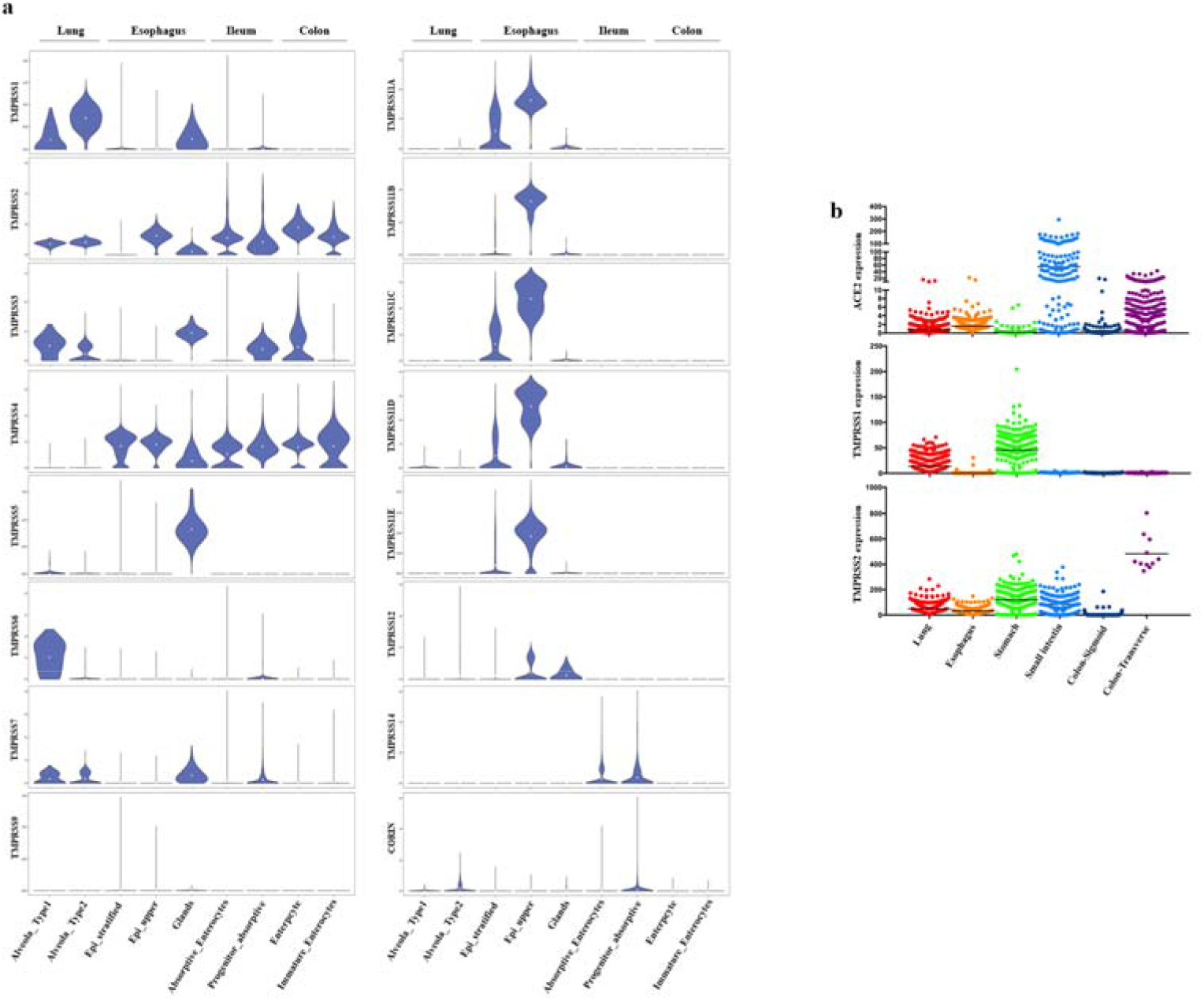
The expression levels of ASE2 and TMPRSS family genes in lung and digestive tracts. a. The violin plots of TMPRSS family genes in lung and digestive tracts. The expression is measured as the mean log2 (TP10K+1) value. b. The expression levels of ACE2, TMPRSS1 and TMPRSS2 verified by RNA-seq data from the GTEx database.

